# Intrinsically disordered regions regulate both catalytic and non-catalytic activities of the MutL*α* mismatch repair complex

**DOI:** 10.1101/475152

**Authors:** Yoori Kim, Christopher M. Furman, Carol M. Manhart, Eric Alani, Ilya J. Finkelstein

## Abstract

Intrinsically disordered regions (IDRs) are present in at least 30% of the eukaryotic proteome and are enriched in chromatin-associated proteins. Using a combination of genetics, biochemistry, and single-molecule biophysics, we characterize how IDRs regulate the functions of the yeast MutL*α* (Mlh1-Pms1) mismatch repair (MMR) complex. Shortening or scrambling the IDRs in both subunits ablates MMR *in vivo*. Mlh1-Pms1 complexes with shorter IDRs that disrupt MMR retain wild-type DNA binding affinity but are impaired for diffusion on both naked and nucleosome-coated DNA. Moreover, the IDRs also regulate the ATP hydrolysis and nuclease activities that are encoded in the structured N- and C-terminal domains of the complex. This combination of phenotypes underlies the catastrophic MMR defect seen with the mutant MutLα *in vivo*. More broadly, this work highlights an unanticipated multi-functional role for IDRs in regulating both facilitated diffusion on chromatin and nucleolytic processing of a DNA substrate.

## Introduction

Intrinsically disordered regions (IDRs) are structurally heterogeneous protein domains that encode diverse functions. IDRs are conformationally flexible, facilitating interactions with multiple partners through intramolecular and intermolecular mechanisms (1, 2). IDRs are often found as linkers connecting functional domains where they can regulate protein stability (1). IDRs are prevalent in chromatin-binding proteins, and the IDRs in these proteins have been implicated in bridging DNA strands, chromatin remodeling, and interacting with other key proteins in DNA metabolic pathways (3, 4). Moreover, IDRs in transcription factors and single-strand DNA binding (SSB) proteins have been reported to tune the DNA binding affinities of these proteins (5–10). Whether these IDRs also regulate scanning on chromatin and other catalytic processes is an open question. This is partly because mutations in such regions often do not confer a specific phenotype, and in some cases, the amino acid sequences contained within IDRs, which are typically poorly conserved among family members, can be critical for the function of a specific IDR-containing protein. Using the mismatch repair protein Mlh1-Pms1 as a case study, we explore the role of IDRs in regulating the DNA scanning and enzymatic activities of a critical eukaryotic DNA repair factor.

The MutL homolog family protein MutL*α*(MLH; Mlh1-Pms1 in baker’s yeast) is essential for eukaryotic DNA mismatch repair (MMR). Mlh1-Pms1 organizes into a ring-like structure that links the ordered N- and C-terminal domains via 160-290 amino acid-long IDRs (11–16) (**Figure 1A**; amino acids 335-499 in Mlh1, 364-659 in Pms1). Mlh1-Pms1 searches for MutS homologs (MSH) bound to DNA mismatches (16–18). A latent MLH endonuclease activity then nicks the newly-synthesized DNA strand resulting in excision of the mismatch (19). This activity requires PCNA, and multiple nicks may enhance the excision step of MMR (20–27).

**Figure 1.**
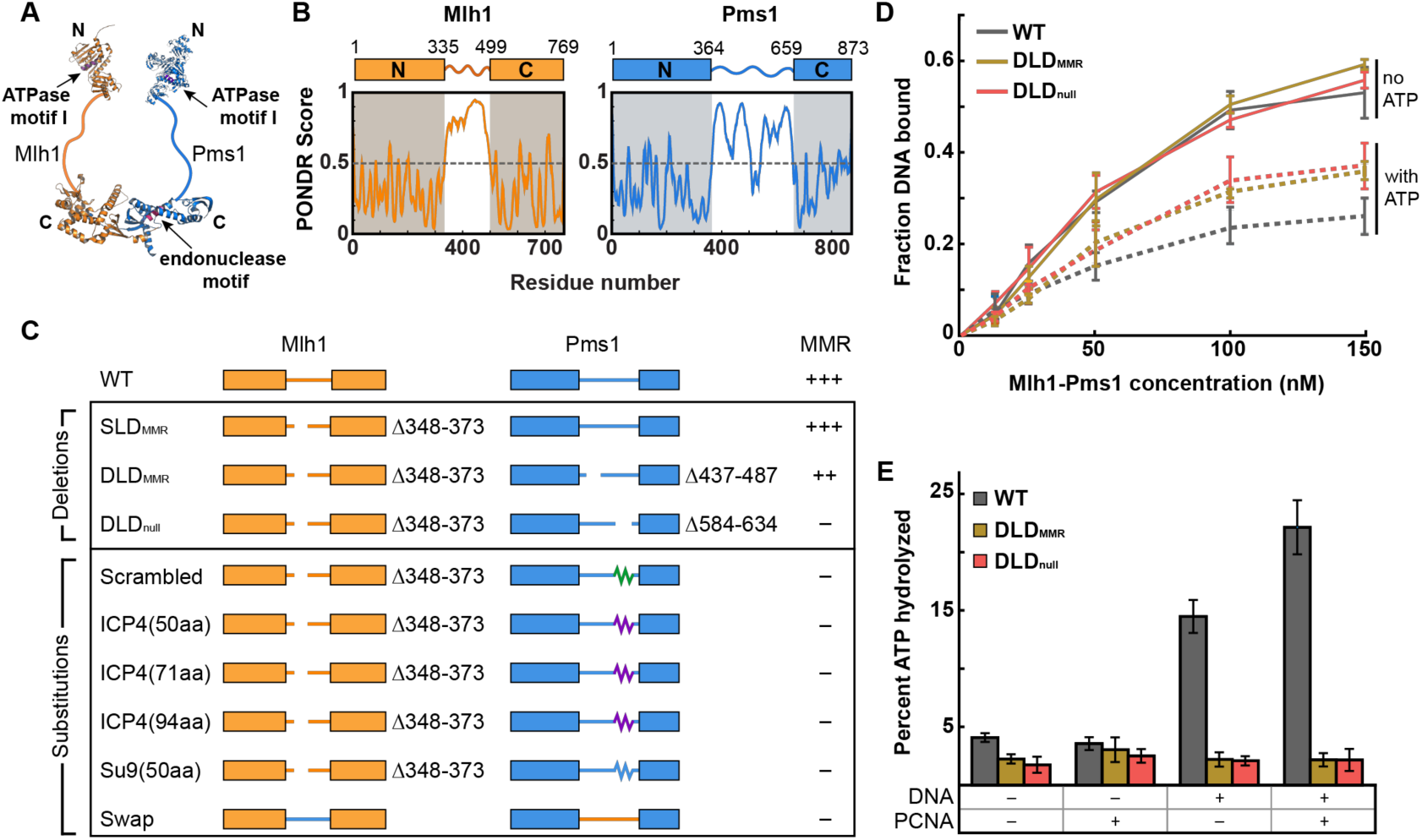
The IDR of Mlh1-Pms1 is critical for MMR *in vivo* and ATP hydrolysis *in vitro*. **(A)** Illustration of Mlh1-Pms1 highlighting the structured N- and C-terminal domains separated by IDRs (solid lines). **(B)** Bioinformatic prediction of long IDRs in both Mlh1 (amino acids 335-499) and Pms1 (amino acids 364-659) using the PONDR VSL2 predictor (73). Any value above 0.5 is considered disordered. **(C)** Schematic of IDR sequence changes made in Mlh1-Pms1, followed by the mutator phenotype conferred by the indicated alleles. +++ wild-type mutation rate, ++ hypomorph, – null. See text for a description of the specific sequences. (D) DNA binding activities for each complex analyzed by filter binding in the presence (dashed line) and absence (solid line) of 1 mM ATP. Mlh1-Pms1 variants were included at final concentrations of 12.5 nM, 25 nM, 50 nM, 100 nM, and 150 nM in buffer containing 25 mM NaCl. DNA binding of a 49 bp oligonucleotide was quantified by scintillation counting. Three replicates were averaged; error bars indicate ± one SD. (E) ATP hydrolysis activities of WT and mutant Mlh1-Pms1 complexes (0.40 μM) were determined alone, and in the presence of PCNA (0.5 μM), or 49-bp homoduplex DNA (0.75 μM), and both PCNA (0.5 μM) and 49-bp homoduplex DNA (0.75 μM). Error bars indicate ± one SD of three replicates.

All MLH-family proteins encode an IDR between the structured N- and C-termini. However, the functional role(s) of the IDRs in Mlh1-Pms1 is enigmatic. The composition and length of the MLH IDRs are critical for efficient MMR in yeast, and missense/deletion mutations within these linkers are found in human cancers (11, 28, 29). We previously proposed that the Mlh1-Pms1 IDRs are sufficiently long to accommodate a nucleosome within the complex, possibly allowing Mlh1-Pms1 to navigate on chromatin *in vivo* (17, 30). In support of this model Mlh1-Pms1 foci that were visualized in live yeast were short-lived (~1.5 min on average), and displayed rapid movements in the nucleus (31). In addition, the Mlh1-Pms1 IDRs display nucleotide-dependent conformational transitions, with ATP binding bringing the N- and C-terminal subunits close together (32–34). This ATPase activity is required for MMR *in vivo* and can stimulate the endonuclease activity *in vitro* (20, 35–38). ATP-dependent conformational rearrangements involving the IDRs are hypothesized to position bound DNA near the endonuclease active site and presumably change Mlh1-Pms1 affinity for DNA (32, 34, 35). Together, these studies suggest that MLH proteins may use conformational changes mediated by the ATP cycle to modulate affinity for DNA, navigate on chromatin, and introduce nicks on a DNA substrate for efficient MMR. However, these possible functions of the IDRs have not been tested directly.

Here, we use a combination of genetics, ensemble biochemistry, and single-molecule biophysics to investigate how the Mlh1-Pms1 IDRs promote both DNA scanning and nuclease activities. We show that both the sequence composition and the precise length of the IDRs are required for optimal MMR *in vivo*. Having mapped genetic requirements for MMR, we next biochemically characterized a double linker deletion (DLD) mutant that was almost completely defective in MMR (DLD_null_), and another linker deletion mutant that retains partial *in vivo* MMR function (DLD_MMR_). Interestingly, both mutants can diffuse on DNA and nick a supercoiled plasmid, but show reduced DNA-dependent ATPase and nucleosome bypass activities. Furthermore, DLDnull is unable to navigate dense nucleosome arrays and is defective in multiple rounds of DNA nicking. These results establish that the IDRs license Mlh1-Pms1 to navigate chromatin and nick DNA at multiple sites to promote efficient MMR *in vivo*. Thus, the IDRs play a critical role in regulating how a DNA repair enzyme scans chromatin for a specific target and how the enzyme activates its endonuclease activity. More broadly, these results expand the functions of IDRs in regulating the DNA scanning and enzymatic activities of chromatin-associated complexes.

## Results

### The IDRs of Mlh1-Pms1 are critical for mismatch repair

We first examined whether the IDRs of Mlh1 (~160 amino acids) and Pms1 (~290 amino acids) contain functionally important amino acids (**Figure 1A–B**). Our previous study established that MMR was ablated in yeast cells that lacked the Mlh1 IDR residues 348-373 and Pms1 residues 548-634 (MMR-null double-linker deletion, DLD_null_; mlh1Δ348-373-pms1Δ584-634). This result was surprising because deleting the same residues in the individual subunits conferred very mild MMR defects (**Supplementary Table S1**) (11). Here, we expand on this early study by defining whether the composition and/or the lengths of the IDRs are critical for supporting MMR.

We first tested whether restoring the IDR of *pms1Δ584-634* to its full length rescued MMR (**Figure 1C, Supplementary Figure S1**). *PMS1* was chosen because truncating its IDR at different positions showed only minor MMR defects and thus may be more likely to restore function with a synthetic linker (11). The substitutions included random scrambling of the 50 critical amino acids (584-634) in Pms1, as well as two biophysically characterized serine-rich regions that were equal or longer than 50 amino acids (obtained from the Herpes Virus ICP4 and *Neurospora crassa* Su9 proteins) (55, 56). All substitutions were initially examined in the wild-type (WT) *MLH1* background, where they did not restore function. The MMR defects conferred by these *pms1* mutants were similar to the *pms1Δ584-634* allele, indicating that the insertions are unlikely to disrupt the stability of the Mlh1-Pms1 complex (**Figure 1C, Supplementary Figure S1**, and **Supplementary Table S1**). If complex stability was compromised, these *pms1* mutants would have shown an MMR defect similar to *pms1Δ* (~8000-fold higher mutation rate compared to *PMS1* in the *lys2::insE-A14* reversion assay, **Supplementary Table S1**). In the *mlh1Δ348-373* background, *PMS1* linker substitutions all conferred a nearly-null MMR phenotype that was reminiscent of the DLD_null_ MMR defect. We also performed a full-length linker swap between the IDRs in *MLH1* and *PMS1* (**Figure 1C, Supplementary Table S1**); these alleles, as swaps or single substitutions, were unable to confer MMR function. Lastly, fine-scale mapping of the *PMS1* 584-634 region using scrambled and single amino acid substitution analyses identified a 20 amino acid region, 594-613,that plays a critical role for the function of the linker. A single substitution in this region, *pms1-Y613A*, conferred a mutator phenotype, (p-value <.00001 to WT; p-value<.00001 to *pms1Δ584-634* compared by Mann-Whitney U test) (**Supplementary Figure S1, Supplementary Table S1**). The *pms1-Y613A* substitution maps to a region of PMS1 that is disordered (**Figure 1B**) and does not encode any known PCNA-interaction motifs, as identified in yeast PMS1 (721QRLIAP), human PMS1 (723QKLIIP), and *B. subtilis* MutL (QEMIVP) (27). Consistent with this, the endonuclease activity of MLH complexes containing the *pms1Δ584-634* mutation is stimulated by PCNA, indicating that this region is not required for PCNA interactions (see below). Together these experiments establish that the specific sequence of the IDR, but not the flexibility, length or disorder is important for efficient MMR.

### IDRs regulate Mlh1-Pms1 ATPase activity in the presence of DNA and PCNA

Mlh1-Pms1 is a DNA-stimulated ATPase and PCNA-activated endonuclease. Nucleolytic cleavage of the newly-synthesized DNA strand by Mlh1-Pms1 is proposed to be a critical strand discrimination signal during MMR (20, 57, 58). We sought to understand the role(s) of the IDRs in promoting the enzymatic activities of Mlh1-Pms1. We compared WT Mlh1-Pms1 to two additional mutant complexes: one mostly functional in MMR (DLD_MMR_; mlh1Δ348-373-pms1Δ437-487), and a second defective (DLD_null_; mlh1Δ348-373-pms1Δ584-634) (**Figure 1**).

All Mlh1-Pms1 variants bound similarly to a 49 bp duplex oligonucleotide in the absence of ATP (**Figure 1D** and **Supplementary Figure S2B**). In the presence of ATP, Mlh1-Pms1 displayed reduced binding to DNA, but both DLD_MMR_ and DLD_null_ displayed DNA binding levels that were higher than WT. These results show that the two DLD complexes are impaired in ATP-dependent interactions with DNA (**Figure 1D**). The ATPase activities of the WT complex are stimulated by DNA and PCNA (27, 34). However, neither DLD complex exhibited such stimulation (**Figure 1E**). We conclude that the IDRs facilitate interactions between Mlh1-Pms1 and DNA, and either directly or indirectly affect the DNA-dependent stimulation of ATP hydrolysis. Remarkably, both DLD_MMR_ and DLDnull showed similar defects in DNA binding and ATPase activities but had very different MMR phenotypes (**Supplementary Table S1**). This puzzle encouraged us to further explore the role of the IDRs in MMR.

### IDRs promote facilitated diffusion on both naked and nucleosome-coated DNA

DNA-binding proteins, including Mlh1-Pms1, locate their targets using facilitated 1-dimensional (1D) diffusion along the genome (17, 18, 30, 59). Based on the biochemical results presented above, we hypothesized that the IDRs of Mlh1-Pms1 are essential for efficient 1D diffusion on chromatin. We examined Mlh1-Pms1 diffusion on double-tethered DNA curtains (**Figure 2A–B**). In this assay, a 48.5 kb-long DNA substrate is extended over a fluid lipid bilayer between two microfabricated chromium barriers (51, 60, 61). The lipid bilayer provides a biomimetic surface that passivates the flowcell surface from non-specific adsorption by DNA-binding proteins. A single FLAG epitope was inserted at amino acid 499 of Mlh1 for downstream fluorescent labeling. The FLAG epitope does not impact Mlh1-Pms1 activities *in vitro* and *in vivo* (17, 30). For fluorescent labeling, Mlh1 was conjugated with an anti-FLAG antibody harboring a fluorescent quantum dot (QD) (17, 30). Using this assay, we characterized WT Mlh1-Pms1, as well as DLD_MMR_ and DLDnull variants. All three Mlh1-Pms1 complexes readily bound DNA and >90% of the molecules rapidly diffused along the entire length of the DNA substrate (**Figure 2C** and **Supplementary Figure S2C**; WT: 97%, N=62/64; DLD_MMR_: 97%, N=79/81; DLD_null_: 90%, N=60/67). Analysis of the movement showed linear mean-squared displacement (MSD) plots, verifying that all three Mlh1-Pms1 complexes freely diffuse on DNA.

**Figure 2.**
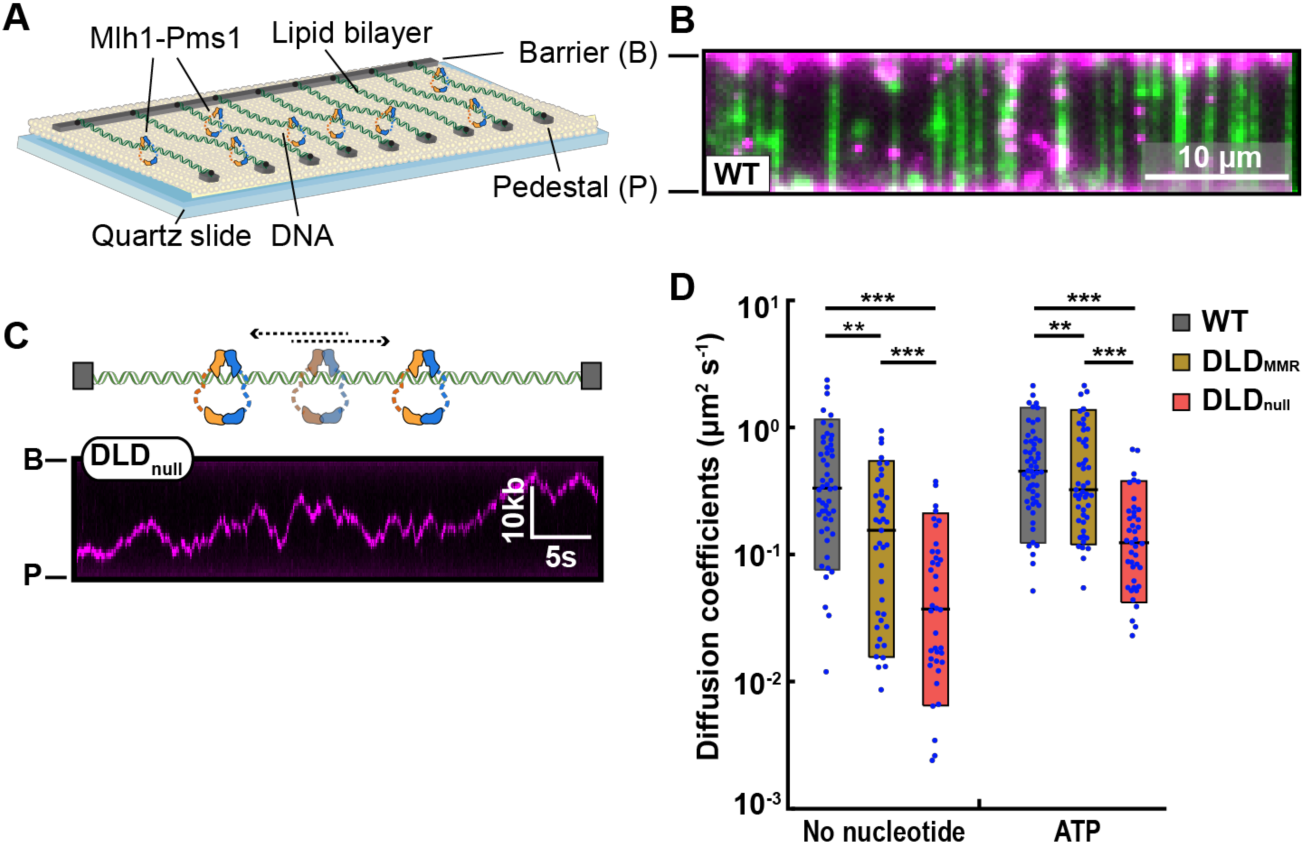
The IDRs promote facilitated Mlh1-Pms1 diffusion on DNA. **(A)** Schematic of the DNA curtains assay. Fluorescently-labeled Mlh1-Pms1 is injected into the flowcell and visualized on double-tethered DNA substrates in the absence of buffer flow. **(B)** An image of Mlh1-Pms1 (magenta puncta) on double-tethered DNA molecules (green). To avoid interference from the DNA-intercalating dye, the DNA is not fluorescently stained during analysis of Mlh1-Pms1 movement on DNA. **(C)** A schematic (top) and representative kymograph of a DLD_null_ Mlh1-Pms1 diffusing on DNA (bottom). **(D)** Diffusion coefficients of Mlh1-Pms1 complexes in the absence and presence of ATP. Boxplots indicate the median, 10th, and 90th percentiles of the distribution. P-values are obtained from K-S test: * *P*-values <0.05, ** *P*-value < 0.01, and *** *P*-value < 0.005.

ATP binding to Mlh1-Pms1 results in dimerization of the N-terminal domains, compaction of the IDRs, and the formation of a ring-like sliding clamp on DNA (32, 34, 36, 62). To probe the functional significance of this conformational change, we measured the diffusion coefficients of the Mlh1-Pms1 variants as a function of ATP. Diffusion coefficients in the ATP-bound state were significantly increased compared to the apo (no nucleotide) condition for all complexes. These results are consistent with a prior single-molecule report of ATP-dependent diffusion of *E. coli* MutL homodimer (59). However, compared to WT and DLD_MMR_, the mean DLDnull diffusion coefficient is ~six-fold lower on DNA in the presence and absence of ATP (**Figure 2D** and **Supplementary Table S2**). While DLDnull displayed the lowest diffusion coefficient of all the complexes in the absence or presence of ATP (**Figure 2D**), DLDnull and DLD_MMR_ displayed similar diffusion coefficients in the presence of ADP or AMP-PNP (**Supplementary Figure S2D**; See Discussion). We conclude that the IDRs of Mlh1-Pms1 are critical for efficient facilitated diffusion on DNA.

Mlh1-Pms1 must efficiently traverse chromatin to locate mismatch-bound MSH complexes. We, therefore, imaged Mlh1-Pms1 on nucleosome-coated DNA substrates. Nucleosomes were assembled using salt gradient dialysis with increasing concentrations of histone octamers to DNA molecules to recapitulate both sparse and dense nucleosome arrays (17, 63). Single nucleosomes were visualized via a fluorescent antibody directed against a triple HA epitope on the N-terminus of H2A. Nucleosomes were distributed over the entire length of the DNA molecule, with a weak preference for GC-rich segments, as described previously (**Supplementary Figure S3A**) (64).

We first determined whether the Mlh1-Pms1 IDRs regulate diffusion past a single nucleosome. DNA substrates with one to seven nucleosomes were assembled into double-tethered DNA curtains (**Supplementary Figure S3B**). Mlh1-Pms1 was added to the flowcell prior to fluorescently labeling the nucleosomes. Keeping the nucleosomes unlabeled guaranteed that Mlh1-Pms1 was not blocked by the H2A-targeting antibody. After recording 10-15 minutes of Mlh1-Pms1 diffusion, a fluorescently labeled anti-HA antibody visualized the nucleosome positions. Diffusing Mlh1-Pms1 complexes encountered and occasionally bypassed individual nucleosomes (**Figure 3A**). To quantitatively determine the probability of bypassing a single nucleosome, we defined a ‘collision zone’ for each nucleosome which encompasses three standard deviations of the spatial resolution of our single-molecule assay (0.08 μm; ~ 300 bp) (**Supplementary Figure S3C** and *Materials and Methods*). Diffusing Mlh1-Pms1 that entered this collision zone from one side of the nucleosome and emerged from the other side was counted as a bypass event. Events where Mlh1-Pms1 entered and emerged from the same side of the nucleosome collision zone were scored as non-bypass encounters. This quantification likely underestimates the frequency of microscopic Mlh1-Pms1 nucleosome bypass events that are below our spatial resolution but does not change any of the underlying conclusions comparing the different complexes.

**Figure 3.**
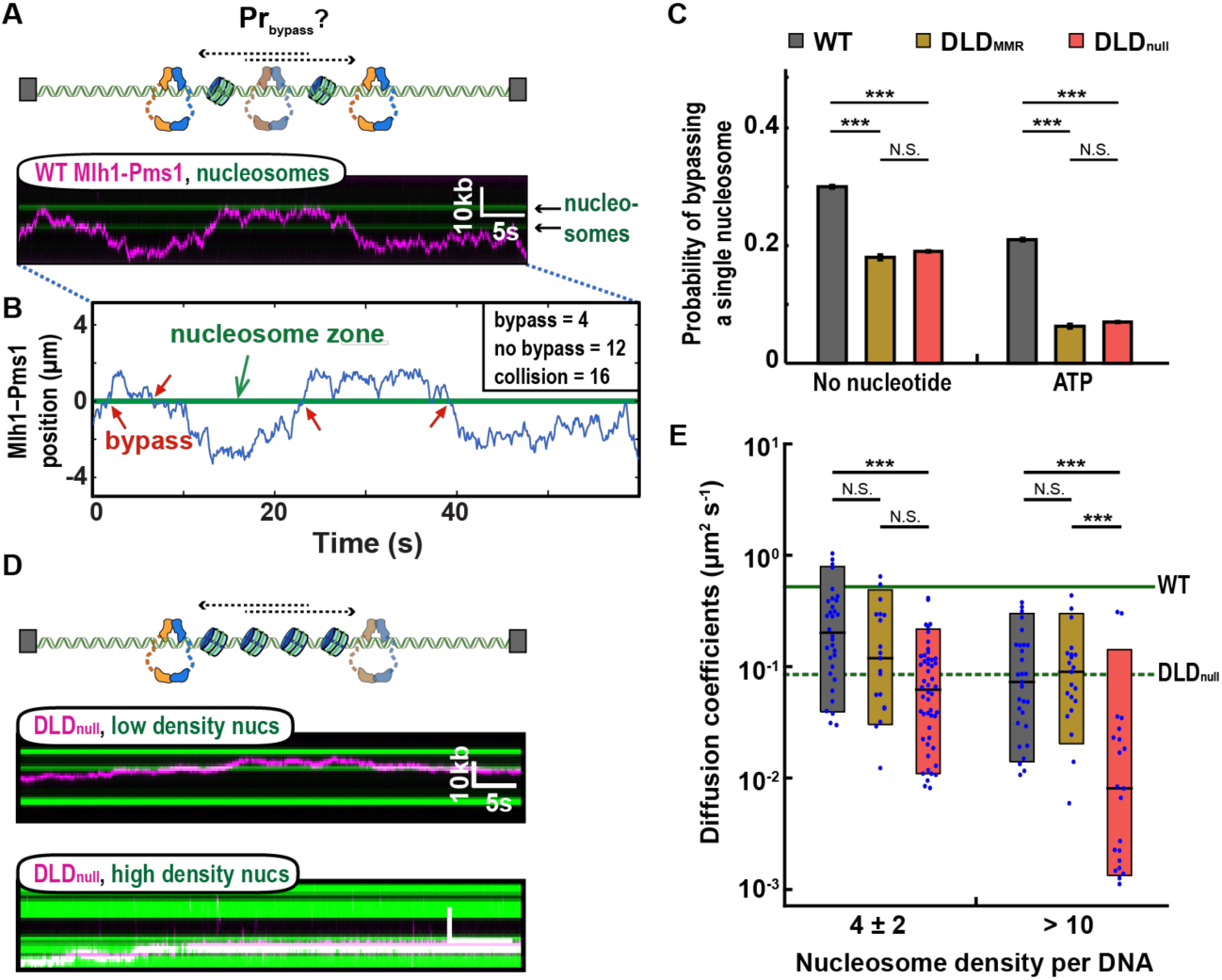
The IDRs increase Mlh1-Pms1 movement on nucleosome-coated DNA. **(A)** An illustration (top) and a representative kymograph of WT Mlh1-Pms1 diffusing past a nucleosome (bottom). **(B)** Trajectory analysis of single nucleosome bypass events. The nucleosome collision zone (green) is defined as three standard deviations of the experimental resolution of the nucleosome position (see *Materials and Methods*). **(C)** The values obtained from the analysis shown in (B) are fit to binary logistic regression to obtain predicted probability of nucleosome bypass. **(D)** A cartoon (top) and representative kymographs of DLD_null_ (magenta) on DNA containing 4 ± 2 nucleosomes (middle, green) or > 10 nucleosomes (bottom, green) in the absence of ATP. Nucleosomes are labeled with fluorescent anti-HA antibodies after Mlh1-Pms1 trajectories are recorded. **(E)** Diffusion coefficients of the three Mlh1-Pms1 complexes on nucleosome-coated DNA. The solid and dashed green lines indicate the mean of the diffusion coefficients of WT and DLD_null_ on naked DNA, respectively. P-values are obtained from K-S test: * *P*-values <0.05, ** *P*-value < 0.01, and *** *P*-value 0.005. N.S. indicates p > 0.05.

WT Mlh1-Pms1 bypassed nucleosomes 30 ± 0.3% of the time (**Supplementary Table S3**). A molecule that travels via a 1D random walk involving facilitated diffusion has a 50% probability of stepping forward or backward on DNA. This 50% probability value is the maximum theoretical bypass probability in the absence of any nucleosome obstacles. Thus, Mlh1-Pms1 is capable of efficiently bypassing a nucleosome obstacle. In contrast, both DLD_MMR_ and DLD_null_ complexes had a 2-fold reduced nucleosome bypass frequency (18 ± 0.5%; N=29 for DLD_MMR_; 19 ± 0.2%; N=27 for DLDnull).

Next, we explored how ATP-induced conformational changes affect nucleosome bypass by Mlh1-Pms1 (**Supplementary Figure 3C**). In the presence of ATP, all Mlh1-Pms1 variants exhibited a reduced bypass probability, with a significantly larger, ~2 to 3-fold, decrease in nucleosome bypass probability for both DLD variants. The decrease in bypass probabilities for DLD_MMR_ and DLDnull mirrors their ATPase activities (**Figure 1E**). Taken together, these data suggest that ATP-dependent dimerization of the N-terminal domains accompanied by conformational compaction of the IDRs reduces dynamic movement on nucleosome-coated DNA.

We reasoned that the combination of a reduced diffusion coefficient and less efficient nucleosome bypass observed with DLD_null_ may compromise its ability to navigate on dense nucleosome arrays. To test this, we increased the histone octamer to DNA ratio during salt dialysis to deposit >10 nucleosomes per DNA substrate (**Supplementary Figure S3D**). At this high density, each nucleosome is optically indistinguishable due to the diffraction limit of light. Nonetheless, by using two-color fluorescent imaging we can still track individual diffusing Mlh1-Pms1 complexes on this nucleosome-coated DNA substrate (**Figure 3D** and **Supplementary Figure S3E**). The 1D diffusion of all Mlh1-Pms1 complexes was restricted on this high nucleosome density substrate compared to naked DNA. Notably, while 1D diffusion coefficients of WT and DLD_MMR_ decreased by 3-fold compared to naked DNA, the DLDnull diffusion coefficient decreased 12-fold on this chromatinized DNA substrate (**Figure 3E**). Thus, the IDRs are important for promoting rapid facilitated diffusion on naked DNA but are especially critical for navigating on chromatin.

### The IDRs are required for multiple rounds of endonucleolytic cleavage

After MSH recognition, Mlh1-Pms1 nicks the mismatch-containing DNA strand for efficient MMR (24, 65). Motivated by the importance of the IDRs in promoting diffusion on both naked and nucleosome-coated DNA, we tested how these domains regulate Mlh1-Pms1 endonuclease activity. We first assayed the ability of Mlh1-Pms1 variants to nick supercoiled DNA in a well-established mismatch- and MSH-independent endonuclease reaction (20, 27, 35, 57). This assay requires the ATP-dependent clamp loader RFC to load PCNA on the closed circle DNA substrate (**Figure 4A**) (66, 67). The endonuclease activity of WT Mlh1-Pms1 was indistinguishable from the DLD_null_ and DLD_MMR_ variants (**Figure 4A** and **Supplementary Figure S4A–B**). However, this assay cannot distinguish between singly- and multiply-nicked DNA substrates. This assay also cannot report the ATP dependence of the Mlh1-Pms1 endonuclease activity because ATP is required for RFC-dependent PCNA loading. To resolve these limitations, we established the alkaline gel-based and single-molecule endonucleolytic assays described below.

**Figure 4.**
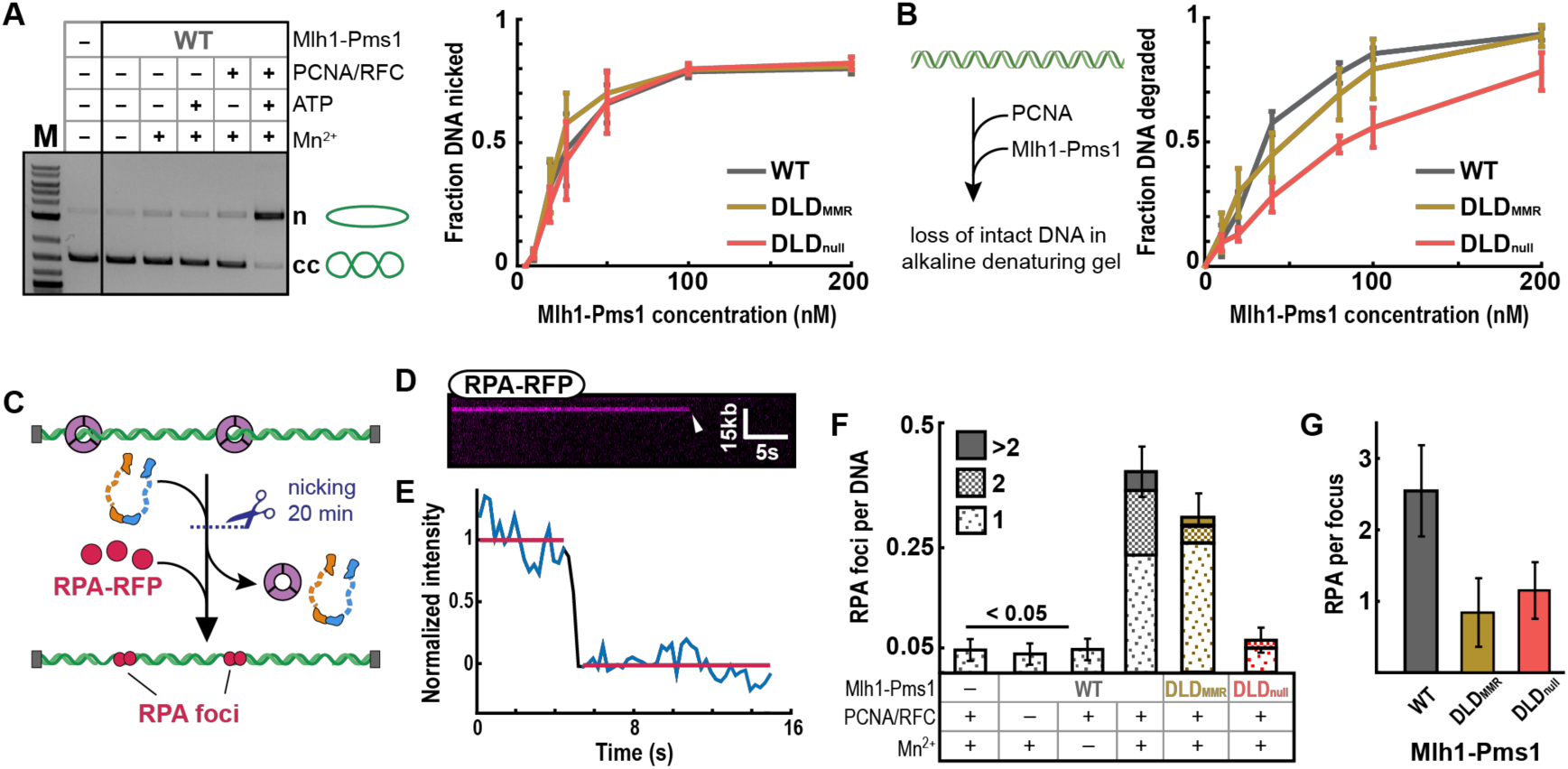
The IDRs regulates extensive DNA nicking. (A) Endonuclease activity on closed circular DNA in the presence (+) or absence (-) of MnSO4, ATP, and yeast PCNA/RFC (left panel). Where + is indicated, the concentration of MnSO4 was 2.5 mM, ATP was 0.5 mM, RFC and PCNA were each 500 nM. The final concentration of WT Mlh1-Pms1 was 100 nM. In the presence of MnSO4, ATP, RFC, and PCNA at the above concentrations, Mlh1-Pms1 variants were titrated from 0-200 nM (right panel). Error-bars: SD of three replicates. (B) Illustration (left panel) and quantification (right panel) of endonuclease activity of wild-type and mutant Mlh1-Pms1 complexes (titrated from 0-200 nM) on linear DNA (also see **Supplementary Figure S4C–D** for controls). All reactions contain 500 nM PCNA, 0.5 mM ATP, and 5 mM MnSO_4_. Error-bars: SD of four replicates. **(C)** Schematic of the single-molecule endonuclease assay. Formation of ssDNA gaps via PCNA-activated Mlh1-Pms1 nuclease activity was visualized by injecting RPA-RFP into the flowcell. **(D)** Kymograph and **(E)** fluorescent intensity profile of an RPA-RFP punctum with a single-step photobleaching event (arrow), indicating a single RPA-RFP molecule on the ssDNA. **(F)** The number of RPA foci per DNA molecule for the indicated Mlh1-Pms1 variants. (G) The number of RPA molecule per punctum for the three Mlh1-Pms1 complexes. To estimate the number of RPA molecules per ssDNA segment, the fluorescent intensity for each punctum was measured and normalized to that of a single RPA-RFP (see *Methods* for details).

We directly tested the role(s) of ATP in Mlh1-Pms1 endonuclease activation on linear DNA substrates analyzed by denaturing gel electrophoresis. PCNA can thread onto the ends of linear DNA, abrogating the need for RFC and ATP (**Supplementary Figure S4C–D**) (27, 57, 68). We observed that Mlh1-Pms1 endonuclease requires PCNA and is further enhanced by ATP binding (**Supplementary Figure S4C–D**). ATP hydrolysis was not required because ATPγS could support the reaction to the same extent or better than ATP, as suggested for the *E. coli* and *Bacillus* MutL (34, 35, 62). Although the DLD_MMR_ variant hydrolyzed linear DNA to approximately the same extent as wild-type Mlh1-Pms1, DNA degradation was attenuated with the DLD_null_ variant (**Figure 4B** and **Supplementary Figure S4D**). This was seen in the presence of ATP, but less so in the presence of ATPγS (**Supplementary Figure S4C**, *see Discussion*). In the assay in Figure 4B, extensive nicking on each DNA molecule accounted for the observed substrate loss, and the reduced nicking by the DLD_null_ complex suggested another *in vivo* MMR defect.

Next, we developed a single-molecule assay to probe the limited nicking that likely occurs for DLD_null_ *in vivo*. This reaction was carried out in two steps. First, PCNA was loaded by RFC on double-tethered DNA curtains in the presence of ATP, as described previously (54). After flushing out RFC, Mlh1-Pms1 was incubated in the flowcell for 20 minutes (**Figure 4D**). PCNA and Mlh1-Pms1 were washed out by 1 M NaCl followed by injecting 50 nM RPA-RFP to visualize the ssDNA gaps made by multiple rounds of Mlh1-Pms1 endonuclease activity (**Figure 4E**). These ssDNA gaps arise from loss of short oligos formed by multiple nicks that are deposited in close proximity by multiple Mlh1-Pms1 molecules. Closely-spaced nicks allow fraying of ssDNAs that are subsequently bound and displaced by RPA (69). Note that we would not be able to detect RPA foci if the nicks on the same strand created by Mlh1-Pms1 were far apart. We quantified the number of RPA foci per DNA and the number of RPA per focus via single-molecule photobleaching. RPA preferentially binds ~30 nt of ssDNA, but individual RPA molecules can bind ssDNA as short as 10 nucleotides (70, 71). Thus, we estimate that puncta with one RPA contain approximately 10-30 nt of ssDNA, whereas puncta with three or more RPA expose > 60 nt of ssDNA. Interestingly, DLD_null_ generated 6-fold fewer RPA foci (0.07 ± 0.02 RPA/DNA; N=307) than WT Mlh1-Pms1 (0.40 ± 0.02 RPA/DNA; N=382). In contrast, DLD_MMR_ was only mildly compromised (0.28 ± 0.02 RPA foci/DNA; N=420) compared to WT complex (**Figure 4F**). We also estimated the length of the exposed ssDNA by counting the number of RPA molecules bound on DNA. The number of RPA per focus was comparable for DLD_null_ (1.1 ± 0.56 RPA; N=20) and DLD_MMR_ (0.9 ± 0.58 RPA; N=68) but was substantially lower than WT Mlh1-Pms1 (2.6 ± 1.2 RPA; N=79) (**Figure 4G**). These data indicate that IDRs are crucial for multiple rounds of DNA nicking during strand excision.

## Discussion

All MLH proteins—from the *E. coli* MutL to the human Mlh1-Pms1— contain IDRs that link the structured N- and C-terminal domains. The importance of these IDRs have been recognized in both bacterial and eukaryotic MMR, but the functions of this domain have remained elusive (11, 16). Here, we show that shortening, scrambling, lengthening, or swapping the IDRs caused mild to severe MMR defects, and even a single amino acid substitution in the IDR of Pms1, Y613A, caused an MMR defect *in vivo* (**Figure 1** and **Supplementary Figure S1**). We, therefore, used three representative Mlh1-Pms1 complexes (WT, DLD_MMR_, DLD_null_) to further probe the mechanistic implications of altered IDRs.

The Mlh1-Pms1 IDRs undergo conformational changes throughout the ATP hydrolysis cycle (32–34, 62). Upon ATP binding, Mlh1-Pms1 adopts a ring-like, scrunched conformation (32). ATP hydrolysis reverts the complex back to the extended open state where it is likely to dissociate from DNA (32, 34, 36, 62). Here, we show that the ATPase activity is disrupted when the IDRs are shortened (**Figure 1E**), indicating that disrupting this conformational cycle feeds back on the ATPase activity encoded in the structured N-terminus of both subunits. These data motivated us to assay the roles of the IDRs in both facilitated diffusion and nucleolytically processing of the DNA.

Mispair recognition by an MSH complex catalytically loads MLH proteins onto DNA. Evidence for Mlh1-Pms1 loading includes an accumulation of Pms1 foci under conditions requiring Msh2-Msh6 and mispaired bases and the identification of *msh6* dominant mutations that prevent Mlh1-Pms1 recruitment *in vitro* and Pms1 foci formation *in vivo* (31, 72). Therefore, Mlh1-Pms1 complexes must scan the genome for mismatch-bound MSH as nucleosomes are being assembled onto the newly synthesized DNA. Strikingly, the DLD_null_ complex is significantly impaired in 1D diffusion on naked DNA and this defect is further exacerbated on dense nucleosome arrays, where the diffusion coefficient of DLD_null_ is decreased by 12-fold compared to that of WT Mlh1-Pms1 (**Figure 2D, 3E**). The different activities of DLD_null_ and DLD_MMR_ suggest that the residues spanning the 584-634 aa region in Pms1 are especially critical for MMR. These residues likely contribute to the conformational rearrangement of the entire complex. A second possibility is that the IDR reorganizes how DNA is channeled through the Mlh1-Pms1 complex. Further structural and biophysical studies will be required to probe the conformational transitions of these IDR variants on DNA. Taken together, our data establish that the IDRs regulate facilitated diffusion of Mlh1-Pms1 on both naked and nucleosome-coated DNA substrates.

Recent studies suggest an alternative EXO1-independent MMR pathway that requires iterative nicking involving multiple Mlh1-Pms1 molecules that are activated via interactions with MSH complexes and PCNA. When Exo1 is absent, multiple nicks may promote strand removal via displacement and/or exonucleolytic activities of Polymerase δ (21, 22, 24, 26, 65). The IDRs may control this activity by ATP-dependent conformational rearrangements that bring the DNA strand close to the nuclease active site. Indeed, ATP-dependent structural rearrangement stimulates the nuclease activity in the bacterial MutL system (35, 62). Consistent with this idea, DLD_null_ was defective in carrying out multiple rounds of DNA cleavage, as seen in both ensemble and single-molecule nuclease assays (**Figure 4**).

**Figure 5** summarizes a working model for how MLH IDRs promote mismatch repair. Mlh1-Pms1 rapidly diffuses on nucleosome-coated DNA in search of lesion-bound MSH complexes. The IDRs play a critical role in promoting facilitated diffusion on chromatin to accelerate the search for MSH-bound lesions. Mlh1-Pms1 is activated by PCNA to nick DNA proximal to an MSH-bound mismatch. This activity may be further regulated by conformational changes in the IDRs that are coupled to ATP hydrolysis. The degree of Mlh1-Pms1 nicking *in vivo* may depend on the concentration of complexes in the vicinity of the mismatch, as well as the availability of the Exo1 nuclease. When Exo1 is unavailable, extensive Mlh1-Pms1-induced nicking provides an alternative strand excision pathway. The loss of MMR observed for the DLD_null_ mutant stems from the combination of defects in ATPase, facilitated diffusion on chromatin, and endonuclease activities. A subset of these phenotypes explains the partial MMR defects of the other IDR variants that we assayed genetically (**Figure 1C** and **Supplementary Figure S1**). Additional studies with the fully-reconstituted mismatch-provoked repair system will provide additional insights into how nicking by Mlh1-Pms1 is regulated at the repair site. More broadly, our results highlight that conformational changes in intrinsically disordered linkers can profoundly alter DNA interactions and enzymatic activities of neighboring structured domains. This work adds additional details to the emerging disorder-function paradigm emerging from biophysical studies of intrinsically disordered proteins.

**Figure 5.**
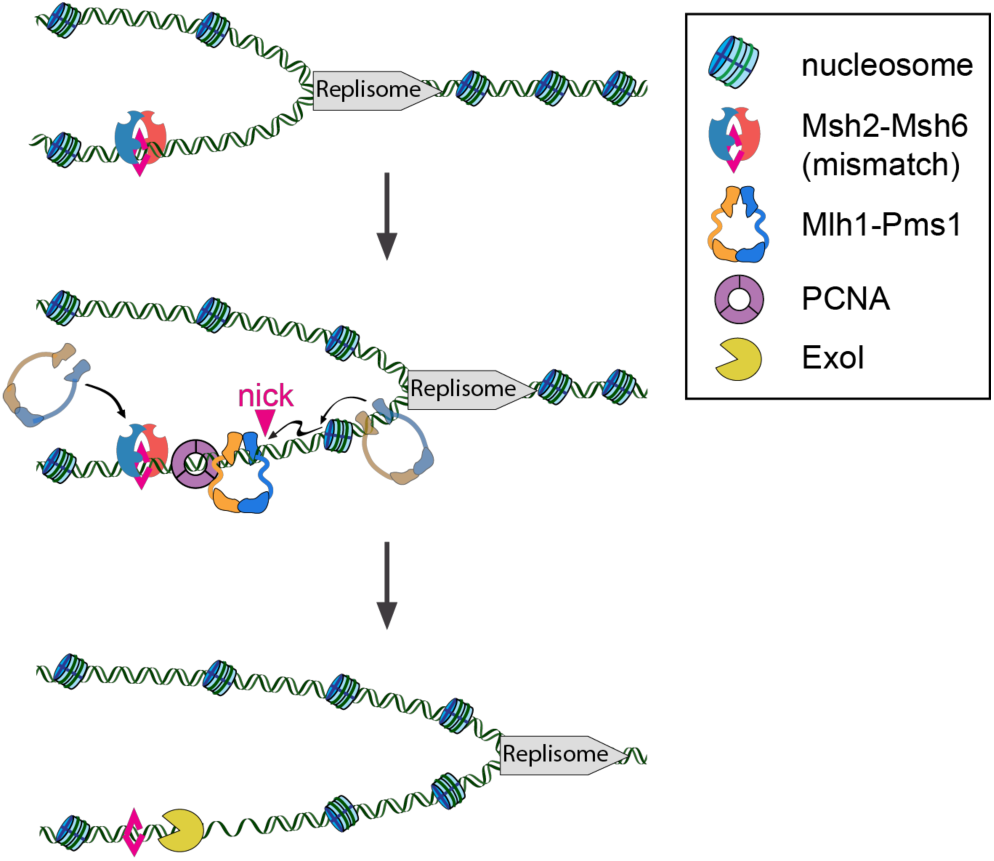
A model of replication-coupled mismatch repair. Msh2-Msh6 locates mismatches in DNA that is nucleosome-free during replication. Mlh1-Pms1 sliding clamps efficiently bypass nucleosomes that are deposited on the newly-replicated DNA. The ability of Mlh1-Pms1 to traverse nucleosomes is important to locate mismatch-bound Msh2-Msh6 but not during endonucleolytic DNA cleavage. Exo1 binds one or more Mlh1-Pms1-generated nicks and degrades the nascent daughter DNA strand.

## Materials and Methods

### Bulk biochemical assays

DNA substrates for bulk biochemical assays: pUC18 (2.7 kb, Invitrogen) was used as the closed circular substrate for endonuclease assays presented in **Figure 4A** and **Supplementary Figure S4A–B**. A 49-mer homoduplex DNA substrate was used in the DNA binding and ATPase experiments presented in **Figure 1D, E, and Supplementary Figure S2B**. This substrate was made as follows. AO3142-5’-GGGTCAACGTGGGCAAAGATGTCCTAGCAAGTCAGAATTCGGTAGC GTG-3’ was labeled on the 5’ end with ^32^P labeled phosphate using T4 polynucleotide kinase (New England Biolabs). Unincorporated nucleotide was removed using a P30 spin column (BioRad). The two oligonucleotides were annealed by combining end-labeled AO3142 with a 2-fold molar excess of unlabeled AO3144-5’-CACGCTACCGAATTCTGACTTGCTAGGACATCTTTGCCCACGTTGACCC-3’ in buffer containing 10 mM Tris-HCl, pH 7.5, 100 mM NaCl, 10 mM MgCl_2_, and 0.1 mM EDTA. Annealing was accomplished by incubating the DNA substrates at 95 °C for 5 min, followed by cooling to 25 °C at a rate of 1 °C/min. Following annealing, excess single-stranded DNA was removed using an S300 spin column (GE). 2.7 kb pUC18 for endonuclease assays on circular DNA was purchased from Thermo. For **Figure 4B** and **Supplementary Figure S4C–D**, the pBR322 plasmid (4.4 kb, Thermo) was linearized using *Hind*III (NEB) by incubation at 37 ºC for 60 min, followed by enzyme inactivation at 80 ºC for 20 min. Linearized fragments were isolated using a PCR clean-up kit (Zymo Research).

#### Protein purification

Yeast WT, DLD_MMR_ and DLD_null_ Mlh1-Pms1 variants (**Supplementary Figure S2A**) were purified from galactose-induced *S. cerevisiae* BJ2168 (*MATa, ura3–52, leu2–3, 112, trp1–289, prb1–1122, prc1–407, pep4–3*) containing expression vectors as previously described (11, 39). Mlh1 contains a FLAG tag at position 499 in wild-type at the equivalent position in Mlh1 truncation mutants. Yeast RFC and PCNA were expressed and purified from *E. coli* (40, 41). RPA-RFP was expressed and purified from Rosetta(DE3)/pLysS cells as described previously (42).

#### Endonuclease assay

Endonuclease reactions were performed in a buffer containing: 20 mM HEPES-KOH (pH 7.5), 20 mM KCl, 2.5 mM MnSO_4_, 0.2 mg/mL BSA, and 1 % glycerol(43). Reactions were stopped by the addition of 0.1 % SDS, 14 mM EDTA, and 0.1 mg/mL Proteinase K(NEB). For reactions on a circular DNA substrate, products were resolved by 1.2 % agarose gel containing 0.1 μg mL^-1^ ethidium bromide, which causes covalently closed circular DNA isoforms to separate from nicked DNA product. Gels were run in 1x TAE (Tris-acetate-EDTA) at 100 V for 45 min. Negative control lanes were used as background and were subtracted out of reported quantifications. Endonuclease assays on linear substrates were carried out and stopped as described for circular DNA substrates. Denaturing agarose gels consist of 1 % (w/v) agarose, 30 mM NaCl, 2 mM EDTA pH 7.5 run in a buffer containing 30 mM NaOH and 2 mM EDTA(44). Immediately prior to sample loading, reactions were supplemented with 30 mM NaOH, 1 mM EDTA, 3 % glycerol, and 0.02 % bromophenol blue (final concentrations), heated for 5 min at 70 ºC, then cooled for 3 min on ice. Gels were run at 50 V for ~3 h. After running, alkaline agarose gels were neutralized in 0.5 M Tris base (pH 7.5) for 30 min and stained with 0.5 μg mL^-1^ ethidium bromide for ~2 h. GelEval (FrogDance Software, v1.37) was used to quantify gels.

#### Filter binding assay

DNA binding assays were performed as described previously (45). Briefly, 20 μL reactions containing 4 nM ^32^P-labeled homoduplex substrate and 11 nM unlabeled homoduplex substrate were combined with increasing amounts of protein in a reaction buffer containing 20 mM Tris-HCl (pH 7.5), 20mM NaCl, 0.01 mM EDTA, 2 mM MgCl_2_, 40 μg mL-1 BSA, and 0.1 mM DTT. Assays with nucleotide contain 1 mM ATP. Reactions were incubated for 10 min at 30 ºC after addition of WT, DLD_MMR_ and DLD_null_ Mlh1-Pms1. Reactions were then filtered through KOH-treated nitrocellulose filters using a Hoefer FH225V filtration device for approximately 1 min. Filters were analyzed by scintillation counting to determine DNA binding efficiency.

#### ATPase assay

ATPase activity was determined using the Norit A absorption method as described previously (43). Briefly, 30 μL reactions contained 0.4 μM of Mlh1-Pms1 (WT, DLD_MMR_ and DLD_null_), 100 μM [γ-32P]-ATP, 20 mM Tris, pH 7.5, 2.0 mM MgCl_2_, 0.1 mM DTT, 1 mM MnSO_4_, 75 mM NaCl, 1 % glycerol, 40 μg/ml BSA. Reactions were incubated for 40 min at 37 °C. When specified, DNA (49-mer homoduplex DNA substrate as described above) and PCNA were included at 0.75 μM and 0.5 μM, respectively.

#### Strains and plasmids

Yeast strains were grown in yeast extract/ peptone/dextrose, minimal complete, or minimal selective media (46). Plasmids used in this study are listed in **Supplementary Table S4**. Full details of plasmid and strain constructions are available upon request. Expression vectors were derived from pMH1 (*GAL1-MLH1-VMA-CBD,2μ, TRP1*) and pMH8 (*GAL10-PMS1,2μ, LEU2*) (39).

#### Linker arm replacement series

A series of *ARS-CEN* vectors were created to test if the 50 amino acid deletion made in the Pms1 linker arm (*pms1Δ584–634*) could be replaced by other sequences (**Supplementary Table S4**). These vectors were derived from pEAA238, which expresses *PMS1* from its native promoter (47). Vectors used to overexpress and purify Mlh1-Pms1 were derived from pMH1 (*GAL1-MLH1-VMA-CBD,2μ, TRP1*) and pMH8 (*GAL10-PMS1,2μ, LEU2*) (39). Insertion plasmids were constructed using NEB HiFi DNA Assembly cloning (pEAA644-656) and Q5 mutagenesis (pEAA659-665). The desired DNA sequence (PCR amplified from specific plasmid or constructed as gBlocks, IDT) was inserted into the deleted region (amino acids 584 to 634) of the Pms1 linker (**Supplementary Table S4**). The DNA sequence of vectors constructed using PCR amplified vector backbones and linker inserts were confirmed by DNA sequencing (Cornell BioResource Center).

### lys2::insE-A14 reversion assay (**Supplementary Table S1**)

Assays were performed as described previously (11). Briefly, pEAA238 (*PMS1*), pEAA548 (*pms1∆584-634*) and derivative linker insertion plasmids of pEAA548 were transformed into EAY3097 (*MATa, ura3–52, leu2Δ1, trp1Δ63, his3Δ200, lys2::insE-A14, pms1Δ::KanMX4*) using standard methods (46, 48). Plasmids were maintained by growing strains in minimal selective histidine dropout media. When tested in combination, pEAA238, pEAA548 (*pms1∆584-634*) or derivative linker insertion plasmids were co-transformed with pEAA213 (*MLH1*) or pEAA526 (*mlh1∆348–373* (FLAG499)) into EAY1365 (*MATa, ura3–52, leu2Δ1, trp1Δ63, his3Δ200, lys2::insE-A14, mlh1Δ::KanMX4, pms1Δ::KanMX4*). Plasmids were maintained by growing strains in minimal selective histidine and leucine dropout media. Null controls were transformed with pRS413 and pRS415 dummy vectors (49). Rates of *lys2::insE-A14* reversion were calculated as μ=f/ln(N·μ), where f is reversion frequency and N is the total number of revertants in the culture (50). For each strain, 15–45 independent cultures, obtained from two to three independent transformants bearing a unique allele, were assayed to determine the mutation rate; 95% confidence intervals and all computer-aided rate calculations were performed as previously described (11).

### Single-molecule experiments and analysis

Data collection on TIRF microscopy: All single-molecule images were collected with Nikon Ti-E microscope equipped with a customized prism-TIRF configuration. The fluorescent samples were illuminated by a 488 nm laser (Coherent) or 532 nm laser (Coherent) through a quartz prism (Tower Optical Co.) depending on the fluorescent dye used. The laser light was adjusted to deliver 40 mW or 15 mW of power at the front face of the prism for 488 nm or 532 nm laser, respectively. Fluorescence was collected by two EM-CCD cameras (Andor iXon DU897, −80°C) using a 638 nm dichroic beam splitter (Chroma), and NIS-Elements software (Nikon) was used to collect the single-molecule data at 50 - 100 ms frame rates. All images were saved as TIFF files without compression for further image analysis in ImageJ (NIH). Experiments were conducted on a floating TMC optical table to avoid spatial drift.

#### Preparation of single-molecule DNA substrates

DNA substrates for single-molecule imaging were prepared by modifying the cohesive ends of λ-DNA (New England Biolabs; NEB). Briefly, 125 μg λ-DNA was mixed with 2 μM IF003 and IF004 in T4 DNA ligase reaction buffer (NEB) and heated to 70°C for 15 minutes followed by gradual cooling to 15°C for 2 hours. After the oligomer hybridization, T4 DNA ligase (2000 units; NEB) was added to the mixture and incubated overnight at room temperature to seal nicks on DNA. The ligase was inactivated with 2 M NaCl, and the reaction was passed over an S-1000 gel filtration column (GE) to remove excess oligonucleotides and proteins. Typically, ~ 10 mL fractions from the first peak were collected and stored at 4°C.

Nucleosomes were deposited on the DNA substrate as described previously with minor modifications (51). The DNA substrate was ligated to the oligo handles, mixed with sodium acetate (pH 5.5) to 0.3 M and isopropanol to 1:1 (v/v), and then precipitated by centrifugation at 15,000 g for 30 minutes. The invisible DNA precipitate was washed with 70% ethanol and dissolved in 2 M TE buffer (10 mM Tris–HCl pH 8.0, 1mM EDTA, 2 M NaCl) to obtain concentrated DNA at ~ 150 ng μL^-1^. For reconstitution, 0.8 nM of the DNA was prepared in 2 M TE buffer with 1 mM DTT for a total volume of 100 μL. Human histone octamers (3xHA H2A with wild-type H2B, H3, H4; Histone Source) were added to the DNA, and the mixture was dialyzed using a mini dialysis button (10 kDa molecular weight cutoff, BioRad) against 400 mL dialysis buffer (10 mM Tris-HCl pH 7.6, 1 mM EDTA, 1 mM DTT, and gradually decreasing concentration of NaCl). The salt gradient dialysis was performed in a cold room and started with 1.5 M NaCl dialysis buffer for 1 hour. The buffer was exchanged every 2 hours to decrease salt in the order of 1 M, 0.8 M, 0.6 M, 0.4 M, and 0.2 M. The last 0.2 M NaCl buffer was used for overnight dialysis. The ratio of DNA to octamer was adjusted to have 3 to 10 nucleosomes per DNA for single nucleosome bypass experiments.

#### Imaging Mlh1-Pms1 on DNA curtains

The Mlh1-Pms1 complexes used in this study contain a FLAG epitope tag at residue 499 on the Mlh1 subunit. We have previously confirmed that placing a FLAG epitope at this position supports full MMR activity *in vivo*, does not disrupt Mlh1-Pms1 biochemical activities (e.g., ATPase, nuclease, MSH2-6 interactions), and is suitable for single-molecule imaging (11, 15, 17, 30). 25 nM of FLAG-tagged proteins were conjugated with 30 nM biotinylated anti-FLAG antibody (Sigma-Aldrich, F9291-2MG) and 25 nM streptavidin QDs (Life Tech, Q10163MP) in a total volume of 60 μL on ice for 7 minutes. The mixture was supplemented with 100 μL biotin and diluted to a total volume of 150 μL in BSA buffer (40 mM Tris-HCl pH 8.0, 1 mM MgCl_2_,0.2 mg mL^-1^ BSA, 50 mM NaCl, 1 mM DTT). The fluorescently labeled proteins were injected into the flowcell immediately after the conjugation at a 200 μL min^-1^ flow rate.

Mlh1-Pms1 loading on DNA is sensitive to the salt concentration in the loading buffer. Therefore, we developed a protocol to efficiently load the fluorescently-labeled protein onto DNA curtains. Mlh1-Pms1 was initially injected into the flowcell containing double-tethered DNA curtains with BSA buffer and 50 mM NaCl to assist its DNA binding. Next, the buffer was switched to imaging buffer (40 mM Tris-HCl pH 8.0, 1 mM MgCl_2_, 0.2 mg mL-1 BSA, 150 mM NaCl, 1 mM DTT, 1 mM nucleotides as indicated). After the flowcell was completely washed with the imaging buffer, flow was terminated to observe 1D diffusion on doubly-tethered DNA substrates. We note that Mlh1-Pms1 diffusion trajectories were indistinguishable between this protocol and complexes that were both loaded and imaged at 150 mM NaCl concentration (**Supplementary Figure S2C**).

#### Fluorescent labeling of nucleosomes

Nucleosomes were fluorescently labeled *in situ* after Mlh1-Pms1 diffusion trajectories were recorded on the DNA substrates. An anti-HA antibody targeting (Immunology Consultants Laboratory, RHGT-45A-Z) was diluted 100-fold in BSA buffer and injected into the flowcell at 10 nM final concentration for 5 minutes. Next, 10 nM secondary antibody was injected and incubated for 7 minutes, then buffer flow was stopped to visualize nucleosomes on double-tethered DNA molecules. We have used anti-rabbit Alexa488 (Life Tech, A-11008) or anti-rabbit ATTO647N (Sigma-Aldrich, 40839-1mL) for the secondary antibody.

#### Particle tracking

Fluorescently-labeled proteins were tracked in ImageJ with a custom-written particle tracking script (available upon request) and the resulting trajectories further analyzed in MATLAB (R2015a, Mathworks). The positions of labeled proteins were determined by fitting every single fluorescent particle to a two-dimensional Gaussian distribution, and the series of time-dependent sub-pixel positions generated each trajectory.

Diffusion coefficients are a measure of a molecule’s movement over an entire trajectory whereas nucleosome bypass is a measure of the local stepping through a nucleosome barrier. Thus, two different approaches were used to calculate these two experimental observables. Mlh1-Pms1 diffusion coefficients were determined by using the trajectories of individual moving molecules on double-tethered DNA curtains in the absence of buffer flow. The one-dimensional (1D) mean squared displacement (MSD) of each particle was determined as a function of the time interval, Δ*t* using the following equation:

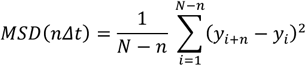

where *N* is the total number of frames in the trajectory, n is the number of frames for a given time interval and ranges from 1 to *N*, Δ*t* is the frame rate, and *yi* is the Mlh1-Pms1 position at frame *i*. The MSD was calculated using the first ten time intervals (e.g. Δ*t* ; 0.05 s to 0.5 s when the frame rate was 0.05 s) and plotted as a function of Δ*t*. Plots were fit to a line and the slope was used to calculate diffusion coefficients of individual Mlh1-Pms1 molecules. Diffusion coefficients were calculated for ≥ 30 molecules in all experiments and are reported as a mean ± standard error of the mean (S.E.M).

#### Measuring single nucleosome bypass frequencies

Fluorescently-labeled Mlh1-Pms1 was loaded onto double-tethered nucleosomal DNA curtains as described above. All nucleosome bypass experiments were done in imaging buffer containing 150 mM NaCl and either no nucleotide or with 1 mM ATP. We determined each collision and bypass event from individual Mlh1-Pms1 trajectories. First, a ‘collision zone’ was defined around each nucleosome position as described in **Supplementary Figure S3C**. Next, the positions of diffusing Mlh1-Pms1 were plotted relative to the center of the nucleosome collision zone. The number of collisions was determined by counting the number of times that Mlh1-Pms1 entered the nucleosome collision zone. Bypass events were defined as collisions that had Mlh1-Pms1 cross from the first to the second side of the nucleosome collision zone. Non-bypass events had Mlh1-Pms1 start and end the collision on the same side relative to the nucleosome. The bypass activity measures how frequently Mlh1-Pms1 passes each nucleosome barrier. To compare the probability of bypassing single roadblock between different conditions with a statistical test, we coded each bypass event as ‘1’ and no bypass as ‘0’ and fit the data to a binary distribution using MATLAB.

#### Statistical methods

We conducted the two-sample Kolmogorov-Smirnov (K-S) test to determine whether average diffusion coefficient differ based on nucleotide types using the PAST3 software package (52). Error bars on the quantified single nucleosome bypass and percentage of moving molecules were calculated in MATLAB using bootstrap analysis with replacement (53). P-values between conditions on single nucleosome bypass experiments were determined in MATLAB using a binary regression model. The significance threshold was set at 0.05 in all tests.

#### Single-molecule nicking assay

5 nM PCNA was loaded by 1.5 nM RFC on double-tethered DNA curtain in Mlh1-Pms1 endonuclease buffer (40 mM Tris-HCl pH 8.0, 0.2 mg mL-1 BSA, 50 mM NaCl, 2 mM MnCl_2_, 1 mM DTT, 1 mM ATP) (54). MgCl_2_ was used instead of MnCl_2_ for manganese negative control. RFC was washed out by injecting endonuclease buffer with 300 mM NaCl for 2 minutes. 20 nM Mlh1-Pms1 complexes were loaded on the PCNA-containing DNA and incubated for 20 min at 30° C followed by washing with 1 M NaCl for 2 min. 50 nM RPA-RFP was then injected to label any gaps larger than 10 nucleotides. For a photobleaching experiment, RPA-RFP was imaged by a 532 nm laser (100 mW at the prism face) with 250 ms exposure time (**Figure 4D–E**). To assess RPA foci, data were collected every 5 seconds with a shutter to reduce photobleaching (**Figure 4F–G**).

## Author contributions

Y.K., C.M.F., and C.M.M. designed and performed experiments. Y.K. performed and analyzed single-molecule experiments. C.M.F. performed genetic assays, ATPase and filter binding assays. C.M.M. performed gel-based nuclease assays. All authors contributed to writing the manuscript.

## The authors declare no conflict of interest

This article contains supporting information online.

### Acknowledgements

We thank Dr. Alba Guarne for sharing information on IDRs and the MLH nicking assay, Dr. Titia Sixma for sharing the MutL over-expression vector, Dr. Andreas Matouschek for sharing plasmids encoding the flexible protein linkers used in the genetic assays, Jeff Schaub for RPA-RFP, Dr. Eric Greene, and members of the Finkelstein and Alani labs for helpful discussions and comments on the manuscript.

